# Identification and Characterization of Calcium Binding Protein, Spermatid Associated 1 (CABS1) in Selected Human Tissues and Fluids

**DOI:** 10.1101/2023.07.21.550040

**Authors:** E. Reyes-Serratos, J. Ramielle L. Santos, L. Puttagunta, S. Lewis, M. Watanabe, A. Gonshor, R Buck, A.D. Befus, M. Marcet-Palacios

## Abstract

Calcium binding protein, spermatid associated 1 (CABS1) is a protein most widely studied in spermatogenesis. However, mRNA for CABS1 has been found in numerous tissues, albeit with little information about the protein. Previously, we identified CABS1 mRNA and protein in human salivary glands and provided evidence that in humans CABS1 contains a heptapeptide near its carboxyl terminus that has anti-inflammatory activities. Moreover, levels of an immunoreactive form of CABS1 were elevated in psychological stress. To more fully characterize human CABS1 we developed additional polyclonal and monoclonal antibodies to different sections of the protein and used these antibodies to characterize CABS1 in an overexpression cell lysate, human salivary glands, saliva, serum and testes using western blot, immunohistochemistry and bioinformatics approaches exploiting the Gene Expression Omnibus (GEO) database. CABS1 appears to have multiple molecular weight forms, consistent with its recognition as a structurally disordered protein, a protein with structural plasticity. Interestingly, in human testes, its cellular distribution differs from that in rodents and pigs, and includes Leydig cells, primary spermatogonia, Sertoli cells and developing spermatocytes and spermatids, Geodata suggests that CABS1 is much more widely distributed than previously recognized, including in the urogenital, gastrointestinal and respiratory tracts, as well as in the nervous system, immune system and other tissues. Much remains to be learned about this intriguing protein.

## Introduction

Using an experimental model in rats to investigate the neural control of inflammatory reactions, we identified a superior sympathetic-submandibular gland axis with anti-inflammatory activity [1–4]. Ultimately, we established that the protein SMR1 (Submandibular rat 1) contains a seven amino acid peptide (TDIFEGG) near its carboxyl terminus with anti-inflammatory activity [3]. Previous work had determined that SMR1 expression was sexually dimorphic, much more abundant in male rats than females, and present in the submandibular glands and testes [5]. SMR1 was predicted to be a prohormone with several polypeptide derivatives with multiple functions [6–11], including vasculature/erectile function, analgesia, and mineralocorticoid-like activity. Our identification of the anti-inflammatory activity was novel, and we characterized its activity in several model systems, including immediate hypersensitivity [12], endotoxic shock [13], asthma [14, 15], and spinal cord injury [16].

However, an analysis of the human genome showed that SMR1 was absent [17], and thus to determine if the anti-inflammatory axis identified in the rat existed in humans, we tested if any human protein contained a heptamer with a similar sequence to that of TDIFEGG in the rat [4, 17, 18]. Human calcium-binding protein, spermatid-associated 1 (hCABS1), was identified given a heptamer TDIFELL near its carboxyl terminus, which was also shown to have anti-inflammatory activity [18]. Moreover, as with SMR1, CABS1 was expressed in testes [19–21], and we identified it in submandibular glands as well [18]. CABS1 was found on the human chromosome analogous to rat chromosome 14, namely chromosome 4, in a region sharing several analogous genes to those in the rat [17].

We collaborated with an experimental psychologist, Thomas Ritz, to determine if human CABS1 (hCABS1) was under autonomic control like SMR1. We established that levels of immunogens in saliva that reacted with anti-hCABS1 polyclonal antibody (pAb) were associated with acute experimental stress [22]. To extend these studies, we developed additional polyclonal and monoclonal antibodies (mAb) to hCABS1. The current study characterizes these antibodies using extracts of testes and submandibular glands, and saliva, serum and hCABS1 overexpression lysate from cultured cells. Our results were then validated using immunohistochemistry to localize hCABS1 in selected tissues, and bioinformatics analyses of CABS1 mRNA expression from public data (Geodata).

## Materials and Methods

### Polyclonal antibodies

Rabbit pAb were made (GenScript Biotech, Piscataway, NJ, USA) against two regions of CABS1 (Fig 1). Polyclonal antibody H1 was raised against hCABS1 amino acid (aa) 375-388 TSTTE<u>TDIFELL</u>KE (underlined anti-inflammatory sequence [18]). Polyclonal antibodies H2.0, H2.1, and H2.2 were raised against hCABS1 aa 184-197 DEADMSNYNSSIKS (region deemed to be highly immunogenic for B lymphocytes) For western blot studies (WB) using pAbs, controls included: (a) pre-immunization serum from each rabbit used to produce each pAb, and (b) blocking controls, performed by incubating each pAb with the respective immunizing peptide in 10X amounts for 18 h (4°C) before WB.

**Figure 1.**
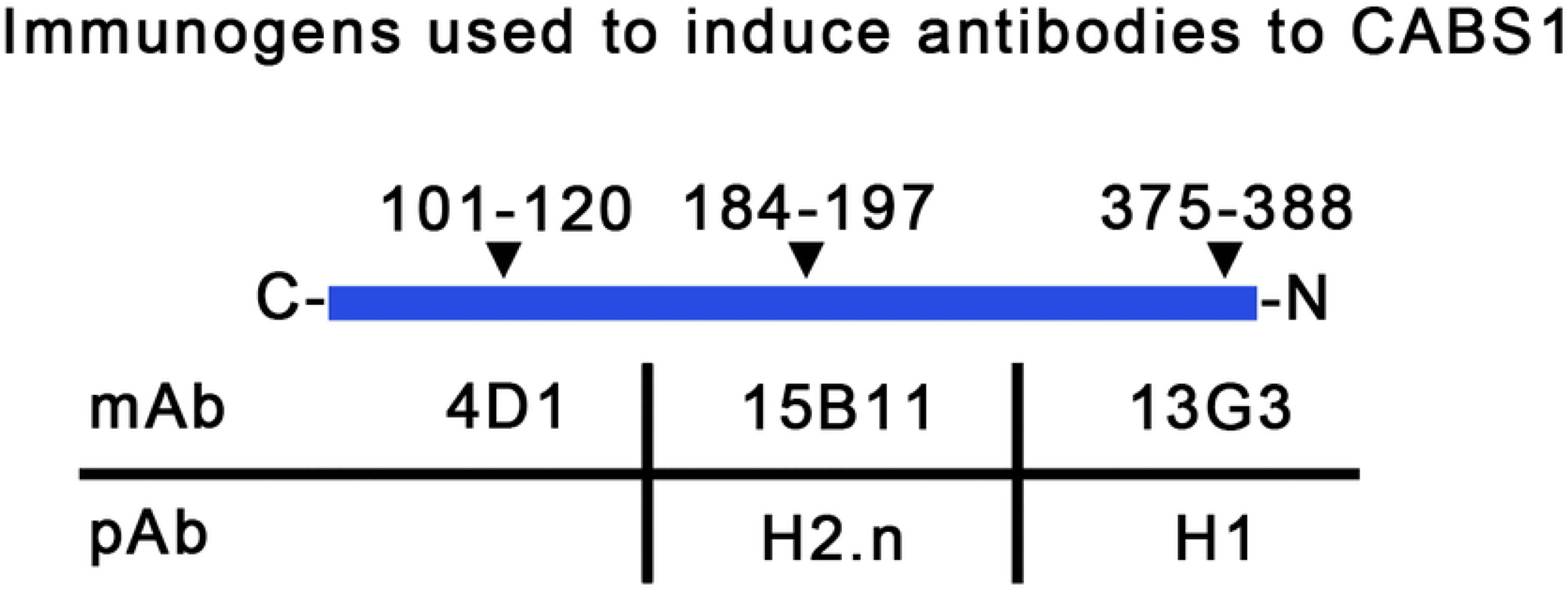
Peptide immunogens used to develop polyclonal (pAb) and monoclonal (mAb) antibodies to different regions of human CABS1. The ranges of the amino acid sequence immunogens are indicated across the top row and the mAb and pAb are listed in the bottom rows. C = carboxyl terminal, N = amino terminal.

### Monoclonal antibodies

Mice were immunized against peptide sequences of hCABS1 (Fig 1). Their spleens were harvested, and spleen-derived B-cells were combined with myelomas to create hybridomas (GenScript Biotech, Piscataway, NJ, USA). Monoclonal antibody 13G3 was raised against hCABS1 aa 375-388 TSTTE<u>TDIFELL</u>KE (underlined anti-inflammatory sequence). Monoclonal antibody 15B11 was raised against CABS1 aa 184-197 DEADMSNYNSSIKS. The immunogen for monoclonal antibody 4D1 was the complete recombinant hCABS1 protein, and the monoclonal was selected to detect a region towards the N terminus (aa 101-120; unpublished observation, R. Buck).

WB controls for mAbs included: (a) no primary antibody, just secondary goat anti-mouse antibodies (LI-COR, Lincoln, NE, USA) or (b) isotype antibodies matched to our mAbs. Mouse IgG1.κ (R&D systems, Minneapolis, MN, USA) was the isotype antibody for mAbs 15B11 and 13G3; mouse IgG2α.κ (Stemcell™ Technologies, Vancouver, BC, Canada) was the isotype antibody for mAb 4D1.

### CABS1 transient overexpression cell lysate

A recombinant hCABS1 overexpression lysate (OEL) produced in Human Embryonic Kidney 293T (HEK293T) cells (OriGene Technologies Inc., Rockville, MD, USA) was used as a positive control in WB. The lysate contains recombinant hCABS1 protein with a FLAG tag (aa sequence: DYKDDDDK) adjacent to its carboxyl end. The negative control cell lysate (NCL) was from HEK293T cells with the same vector but lacking a hCABS1 cDNA insert. As an additional control, we probed the overexpression lysate with ANTI-FLAG ® M2 (Sigma), a mouse monoclonal antibody targeting FLAG sequence (gift from Drs. Steven Willows and Marianna Kulka, Nanotechnology Research Centre, Edmonton, AB).

### Human samples

All tissues and fluids were collected and stored under ethics protocols approved by the University of Alberta Human Ethics Review Board, REB 4 (Biomedical Panel). The Alberta Research Information Services (ARISE) System approved our protocols Pro00001790 and Pro00112432 and provided us with written consent. Recruitment for saliva samples and serum started January 1st 2017 and completed June 30, 2018. Surgical specimens of salivary glands to be used for WB were collected between January 1, 2012 and June 30, 2018. Surgical specimens for IHC studies (secondary use, Pro00112432) were identified (July 30, 2021 to June 30, 2022) from archived samples that were originally collected for clinical diagnostic purposes.

#### Submandibular gland (SMG)

Fresh human SMG samples were collected from patients, female and male, undergoing surgical removal of squamous carcinoma at the University of Alberta Hospital (Edmonton, Canada). A grossly normal looking portion of each SMG was homogenized at 4°C using a diluent RIPA buffer supplemented with P8340 protease inhibitor cocktail (Sigma Aldrich, Markham, ON, Canada). The remaining fresh tissue was frozen (-80°C) for future lysate extraction. Post-homogenization, samples were centrifuged, and the supernatant was collected, aliquoted, and stored at -80°C. Before analyses, each sample’s total protein concentration was determined using a Pierce™ BCA Protein Assay Kit (Thermo Scientific, Waltham, MA, USA).

For immunohistochemical studies, SMG samples removed at surgery for cancers at the University of Alberta Hospital were fixed in 10% buffered formalin. Following paraffin embedding, tissue sections were assessed by microscopy. The sections were stained with anti-hCABS1 antibodies (below), and areas that appeared to be morphologically normal were assessed for the distribution of CABS1 immunoreactivity. One sample of submandibular gland was kindly provided to Dr. Stephen Lewis by GlaxoSmithKline’s (GSK) Immune-inflammation therapy unit’s human biological sample repository, arranged by Dr. Alessandra Giarola of Bioelectronics Research and Development (GSK).

#### Testes

Archived surgical specimens of testis tissue initially collected for clinical pathological analyses were used for immunohistochemical studies; one from a 58 year old with a mass that was determined to be scar tissue, and another from an unknown subject.

#### Blood

Serum was retrieved from a 30-year-old male. A vacutainer tube containing no additive was used to collect 10 mL of whole blood, incubated for 45 min to allow clotting and centrifuged at 1500g, 21°C for 15 min. Serum was aspirated, transferred into 300 µL aliquots, and stored at -80°C.

#### Saliva

Unstimulated whole saliva from a 30 year old male was collected using the passive drool technique protocol (Salimetrics LLC, Carlsbad, CA, USA) [23] over 15 days. Following breakfast and teeth brushing, saliva was collected after 1.5 h. Samples were frozen at -20°C immediately after collection. When collection was finalized, all samples were thawed and pooled. The resulting pool was centrifuged at 1500g, 4°C for 20 min, and the supernatant was transferred into aliquots. Prior to analyses, total protein concentration was determined as above.

### Western blot analysis

For 1 D electrophoresis, we followed standard BIO-RAD protocols. Briefly, the TGX™ FastCast™ 12% Acrylamide Kit (Bio-Rad, Mississauga, ON, CA) was used to prepare 1.5 mm thick gels the day before an experiment and polymerized (30 min) gels were kept at 4°C. On the day of the experiment, 1X WB running buffer (Trizma base – 40 mM, glycine – 300 mM, methanol – 200 mL) was prepared and kept at 4°C.

For WB, electrophoresed samples in gels were placed in a cassette containing fiber pads, filter papers, and a 0.45 µm pore-size nitrocellulose membrane. The cassette was placed in a vertical transfer chamber with an ice pack on the side in WB transfer buffer; transfer was done at 0.5 A for 1 h. The nitrocellulose membrane was washed with double distilled water before adding WB blocking buffer (1% fish skin gelatin [Truin Science Ltd., Edmonton, AB, Canada], 1X PBS, 0.1% Tween20). Blocking was done at room temperature for 1 h. The membrane was probed overnight at 4°C with the relevant titrated antibody(-ies) diluted in a 50:50 solution containing PBS supplemented with 0.05% Tween 20, and blocking buffer. The antibody solution was then removed, and the membrane was washed. LI-COR Biosciences secondary antibodies were diluted 1:10,000 in the same solution as the primary antibodies.

### Molecular weight determination

All WB included a molecular mass reference (M_r_). WB images from studies of pAb produced by LI-COR were opened in Microsoft Paint and the M_r_ of each band (kDa) was recorded. For M_r_ estimates from studies with mAb, calculations were done manually.

### Immunohistochemical analyses

SMG and Testis: Tissue sections on slides were baked at 60°C for 1 h and then deparaffinized and retrieved on Dako Omnis fully automated IHC platform (Agilent, Santa Clara, CA, USA) using EnVision FLEX target retrieval solution, high pH (Dako Omnis) at 97°C for 30 min. Monoclonal antibody 15B11 (targeting the mid region of hCABS1) was diluted 1/100 in EnVision FLEX antibody diluent and was applied for 20 min. at 32°C followed by EnVision Flex+ Mouse LINKER (Dako Omnis) for 10 min. Subsequently, EnVision FLEX, HRP.Rabbit/Mouse was added for 20 min. Visualization was performed using DAB+ substrate chromogen system (Dako Omnis) and counter stained with Gill I Hematoxylin formula (Leica, Wetzlar, Hesse, Germany). All histology and immunostains were reviewed with an Olympus BX41. All histology images were captured with a Nikon Eclipse E600 microscope with an attached Nikon Digital-sight DS-Fi2 Ki7607 camera.

SMG tissue from one 38-year-old female subject (gift from GSK, see above) was sectioned, deparaffinized, rehydrated, and microwaved for 10 min in 10 mM sodium citrate (pH 6) for antigen retrieval (i.e., heat-induced antigen retrieval). Subsequently, sections were blocked for 30 min at room temperature in blocking buffer (PBS containing 5% bovine serum albumin and 0.1% Triton-X 100). Next, blocked sections were incubated overnight at 4°C with the primary pAb (H1, H2.0, H2.1, H2.2, or respective preimmune serum) diluted 1/200 in blocking buffer. The next day, slides were washed with PBS and incubated for 2 h at room temperature with the secondary antibody (goat anti-rabbit IgG H+L conjugated with Alexa Fluor 488, Thermo Scientific), washed with PBS and mounted in VECTASHIELD® HardSet™ antifade medium with DAPI (Vector Laboratories, Burlingame, CA, USA). Images were acquired with a Retiga EXi, Fast 1394 digital camera (Teledyne Photometrics, Tucson, AZ, USA) attached to a Diaphot 200 inverted phase contrast microscope (Nikon) and evaluated using Q-Capture imaging software (Teledyne Photometrics).

### Mass spectrometry sequencing (MS-seq)

Transient overexpression samples, OEL and NCL, were separated by 1-D SDS-PAGE and stained using Blue Silver dye. To assess the distribution and relative abundance of hCABS1, the gels were cut into 11 equal segments and sent for analysis to the Alberta Proteomics and Mass Spectrometry Facility (University of Alberta, Edmonton) for in-gel trypsin digestion and MS-seq analysis using a LTQ Orbitrap XL Hybrid Ion Trap-Orbitrap mass spectrometer (Thermo Fisher Scientific). Data were processed using Proteome Discoverer v.1.4 (Thermo Fisher Scientific) using the Sequest (Thermo Fisher Scientific) database search algorithm; only proteins with ≥2 tryptic peptides identified by MS-seq were included.

### Meta-Analysis of studies from Gene Expression Omnibus (GEO) database

Gene expression data was collected from the GEO database and supplemental data provided in previous studies. Relevant studies were identified using specific keywords, including CABS1, Normal, and Tissue. Non-normal tissue or treated cell/tissue datasets were excluded from the search. Raw data was downloaded and rescaled to 0 to 100% for each dataset using min-max normalization. We combined the data from three studies [24–26] and analyzed the data distribution in a histogram. Over 94% of the data points were observed within 0 to 50%. A continuous graded color scale of the data distribution was used to create the meta-analysis figure.

## Results

### Detection of rhCABS1 in OEL using pAb and mAb

#### pAb

We previously published studies of hCABS1 in OEL using pAb H2.0 [18]. These studies were repeated with pAb H1 and H2.0, H2.1, and H2.2 (Fig 2A). Negative controls using immunogen blocking and preimmune sera from the rabbits used to produce H1 and H2.0, H2.1 and H2.2 were negative. H1, H2.0, and H2.2 specifically detected immunoreactive bands in OEL at 84 and 67 kDa, whereas H2.1 detected the 84 kDa band specifically (Fig 2A; n=8-16 independent experiments). In NCL, these pAbs variably detected bands at approximately 76, 55, 47, 34 and 21 kDa (Fig 2A). Antibody to the FLAG marker of rhCABS1 in OEL detected immunoreactive bands at 84 and 67 kDa; no bands were detected in NCL. Similarly, MS analyses detected hCABS1 in OEL, but not in NCL. CABS1 peptides were detected in 1D gel segments of 119-85, 84-60, 59-40, 39-33, 32-22 and 16-11 kDa (Fig 3), as mentioned briefly in our previous report [18]. Two to 10 unique peptides were identified in the gel segments (Fig 3B) and peptide coverage of the protein in the segments ranged from 7 to 48%. As previously reported [18], there was no evidence that peptides from selected regions of hCABS1 were preferentially found in different segments of the gel. No peptides were detected above 119 kDa, in the 17-21 range or below 10 kDa.

**Figure 2.**
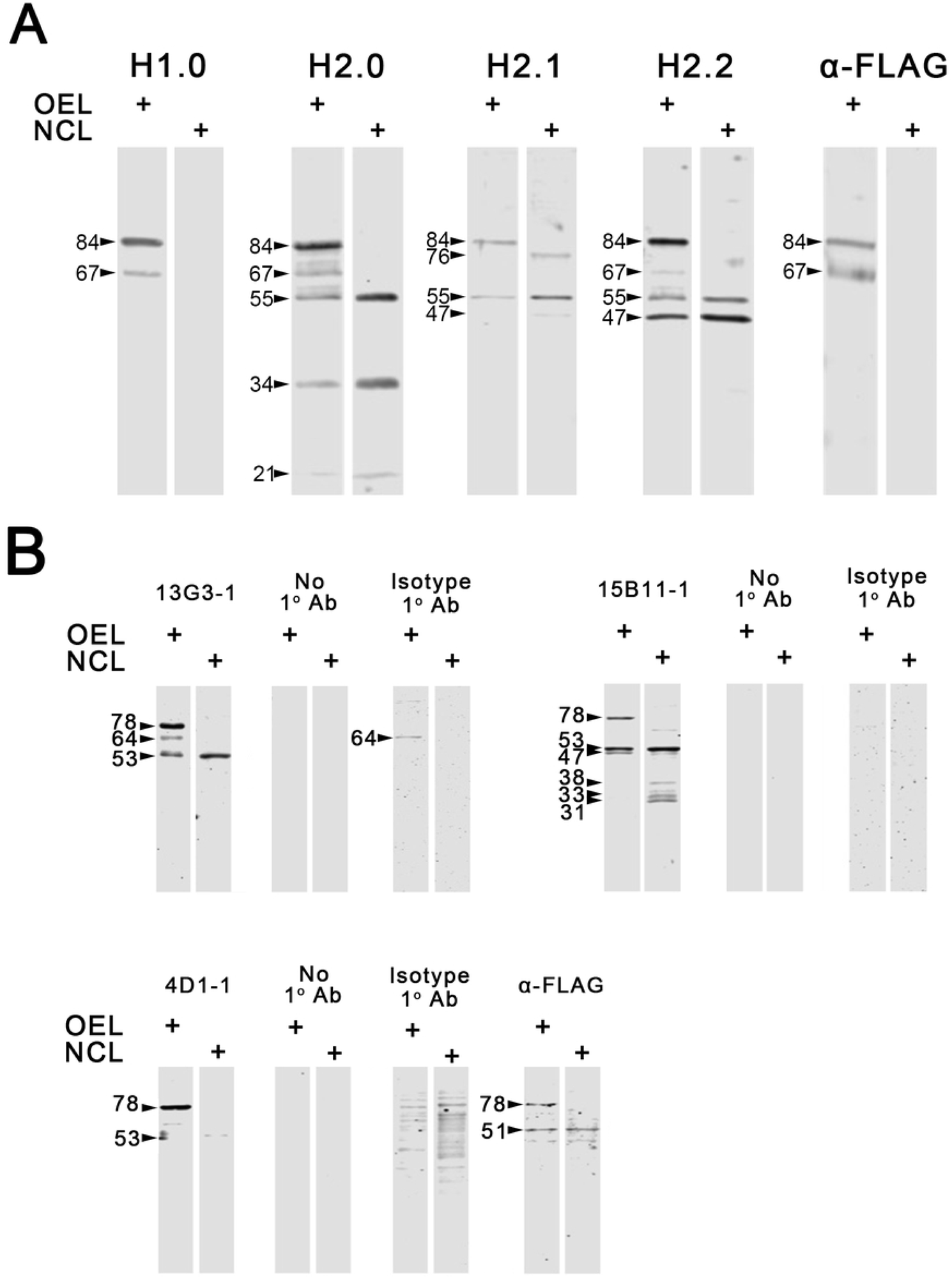
Western blot analyses of human CABS1 in transient overexpression lysate (OEL) of HEK 293T cells and in control lysate (NCL) of HEK 293T cells transfected with empty plasmid. In A, the representative results of four pAb to peptide immunogens of hCABS1 are shown (n = 8 to 16 replicates; 25µg of OEL or NCL lysate were used, for H1 and H2 3 ng/µL and for H2.1 and H2.2 2 ng/µL of pAb used were). Controls with preimmune serum from each rabbit, or with immunogen blocking controls (preincubation with peptide immunogen at 10X concentration of primary antibody for 18 h at 4o C), were negative (not shown). In B, mAb 13G3 (10 ng/µL), 15B11 (1 ng/µL) and 4D1/ (1ng/µL) were used, and 1 µg of OEL or NCL (n =2). Controls with no primary antibody, or with isotype control antibodies are shown. Antibody to the FLAG tag on the rhCABS1 was used as a control for these studies with pAb and mAb.

**Figure 3.**
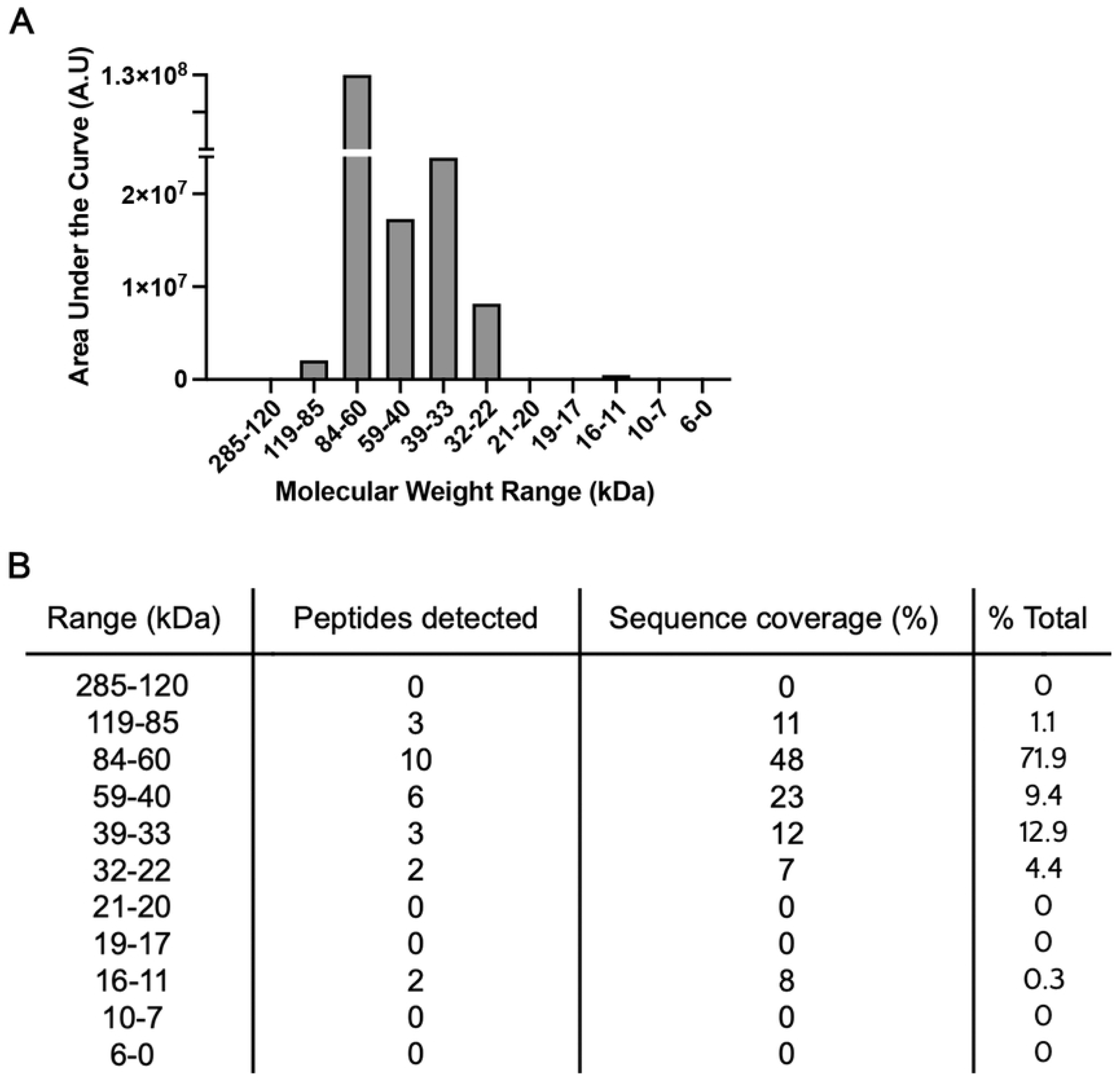
Estimate of the abundance of hrCABS1 in sections (range of kDa) of 1 D gel analyzed by mass spectroscopy. A, relative abundance of rhCABS1 in each gel segment (kDa range); B, the number of rhCABS1 peptides detected in each gel segment, the percentage of rhCABS1 covered by the peptides in each segment and the percentage of the total rhCABS1 recovered.

In summary, our pAbs detected CABS1-specific bands at 84 and 67 kDa in rhCABS1 OEL and not in NCL.

#### mAb: (n=2 independent experiments for each)

Antibody 13G3 (same immunogen as H1 pAb) detected bands at 78, 64, and 53 kDa in OEL, and in NCL, a band at 53 kDa (Fig 2B). No bands were observed in membranes probed with no primary antibody, but OEL probed with isotype antibody IgG1.κ at the same concentration as 13G3 showed a band at 64 kDa (Fig 2B). WB analyses using 15B11 (same immunogen as H2.n pAb) showed immunoreactive bands at 78, 53, and 47 kDa for OEL, and bands at 53, 47, 38, 33, and 31 kDa in NCL (Fig 2B). No immunoreactivity was observed in membranes probed with isotype antibody IgG1.κ at the same working concentration as 15B11 or with no primary antibody. Antibody 4D1 shows a band at 78 kDa in OEL, and a faint band at 53 kDa in NCL (Fig 2B). No signal was detected in membranes probed with no primary antibody, but a membrane probed with isotype antibody IgG2α.κ showed several bands for NCL in a smear pattern in a range between 100 and 35 kDa. Evaluation of these lysates with anti-FLAG antibody showed two bands at 78 and 51 kDa, with the 51 kDa band also present in NCL (Fig 2B).

In summary, our mAbs specifically detected rhCABS1 at 78 kDa in OEL. Given that this band was also FLAG positive, we postulate that the 84 kDa band detected with pAbs and this band detected at 78 kDa with mAbs are the same (see Discussion).

## Detection of hCABS1 using pAbs and mAbs in human fluids and submandibular glands lysate

### hCABS1 in submandibular gland lysate

#### pAb

With each of the four pAb several immunoreactive bands ranging from ∼22 to 127 kDa were detected in male SMG lysate (Fig 4A). Preimmune sera for H1 and H2.0 were used as a negative control and gave no immunoreactive bands (not shown). There were differences among the pAb in the immunoreactive bands that were detected; bands at ∼71, 52 and 27 kDa were detected by at least three of four pAb.

**Figure 4.**
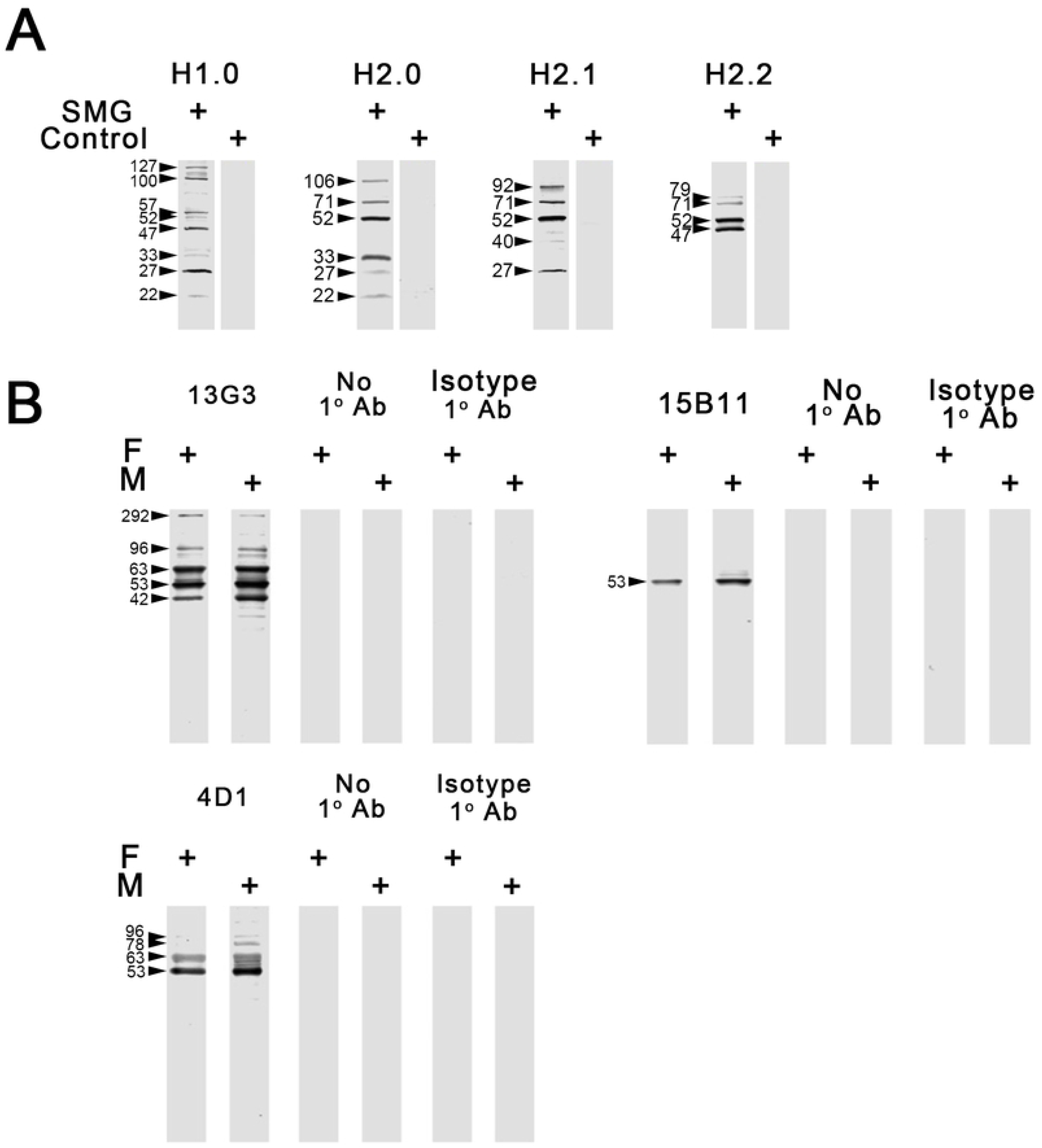
Western blot analyses of CABS1 in human female and male submandibular gland lysates. A) As determined from studies with OEL and NCL (Fig 2), H1 and H2 (3 ng/µL) and H2.1 and 2.2 (2 ng/µL) were used for studies of male SMG lysate (25 µg) (n = 4-5). Immunogen blocking controls (preincubation with peptide immunogen at 10X concentration of primary antibody for 18 h at 4°C), were negative. Preimmune sera controls for H1 and H2.0 were negative also (preimmune sera for H2.1 and 2.2 were not studied. B) Studies with mAb used 10 µg of lysate in each lane; for mAb 13G3 and 4D1 10 ng/µL and for 15B11, 1 ng/µL of mAb was used (n = 3). Controls with no primary mAb or with isotype control mAb are shown and were negative.

#### mAb

No immunoreactive bands were detected when probing membranes with isotype antibody at the same working concentration as the respective primary mAb, or when adding no primary antibody. WB analyses of male and female SMG lysates using 13G3 (n=3) showed immunoreactive bands at ∼96, 78, 63, 53 and 42 kDa. (Fig 4B). WB analyses of SMG lysate using the 15B11 mAb (n=3 independent experiments) detected a single immunoreactive band at 53 kDa in both male and female extracts. WB of SMG lysate using antibody 4D1 (n=3) detected two bands at 63 and 53 kDa in both male and female SMG, and in the male, additional bands at 96 and 78 kDa and between 63 and 53 kDa (Fig 4B).

Thus, with mAbs, SMG lysate has anti-hCABS1 specific immunoreactive bands at ∼96, 78, 63, 53 and 42 kDa with some distinctions among the mAbs used. With pAb, bands at ∼71, 52 and 27 kDa were detected with 3 of 4 pAb.

### hCABS1 in saliva

#### pAbs

Given our experience with careful antigen and antibody titrations using semi-automated capillary nano-immunoassay (NCIA) [23], we reassessed our previous optimization [18] of antibody and antigen concentrations using pAb and human samples. We determined that optimal specificity was achieved at 2.2 µg of human saliva and 0.2 to 0.3 ng/µL of pAbs. At these concentrations in a pilot experiment, no unequivocal bands selective to H1 compared to preimmune serum were detected in saliva, although a band was seen at 60 kDa (Fig 5A). With H2.0 a selective band (versus preimmune serum) was seen at approximately 60 kDa and faint bands were seen at ∼ 90 and 27 kDa (Fig 5A). Studies were not done with H2.1 and 2.2.

**Figure 5.**
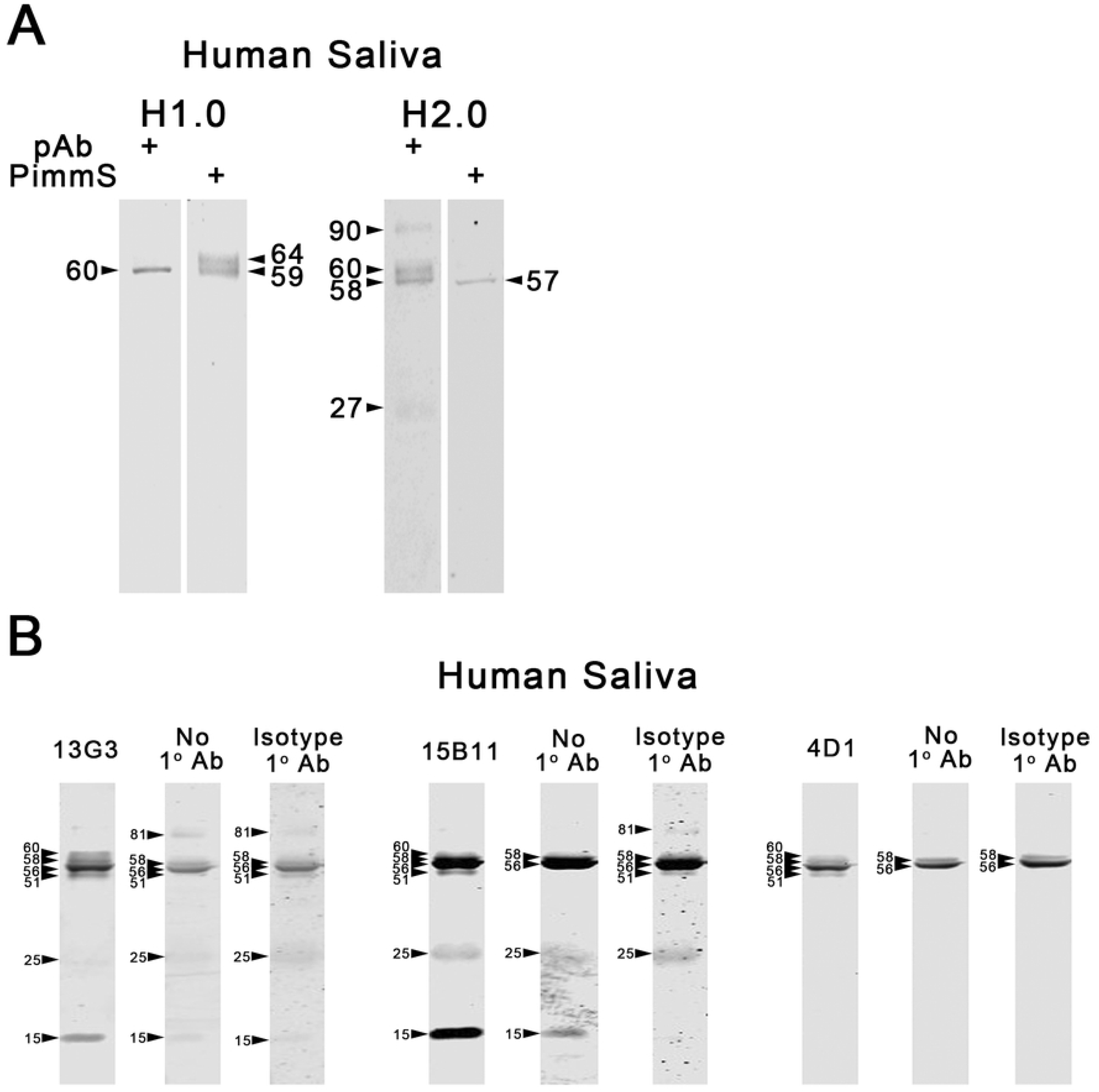
Western blot analyses of CABS1 in human (male) saliva. A) In this experiment (n = 1), optimal specificity with pAb was achieved at 2.2 µg of human saliva and 0.2 to 0.3 ng/µL of pAb and preimmune sera. B) For mAb 13G3 10 ng/µL and for mAb 15B11 and 4D1 1 ng/µL were used together with 10 µg of saliva (n = 3 for each mAb). Controls with no primary mAb or with isotype control mAb are shown.

#### mAbs

As with studies using pAb, studies with all three mAb on salivary supernatants were challenging but appear to detect a specific immunoreactive band of ∼ 60 kDa that was not detected with only secondary antibody or with isotype antibody (Fig 5B).

In summary, results from WB analyses of human saliva with pAb and mAb suggest that there is a specific immunoreactive band of ∼60 kDa (see Discussion).

### hCABS1 in Serum

#### pAbs

For analysis of CABS1 in human serum, we selected 1.5 µg of serum and 0.03 and 0.02 ng/µLof pAbs H.1.0, H2.0 and H2.1, H2.2 respectively as appropriate concentrations of reagents (Fig 6). An initial experiment with H1, 2.0, 2.1 and 2.2 with a human serum sample suggested that CABS1 is present in serum. Several immunoreactive bands were detected with the pAbs that were not detected in the corresponding preimmune sera (Fig 6A). The bands ranged from ∼ 180-135, 90-100, 60–75 and 47-56 kDa. Given this complexity of immunoreactive bands with pAb, we focused studies on mAb.

**Figure 6.**
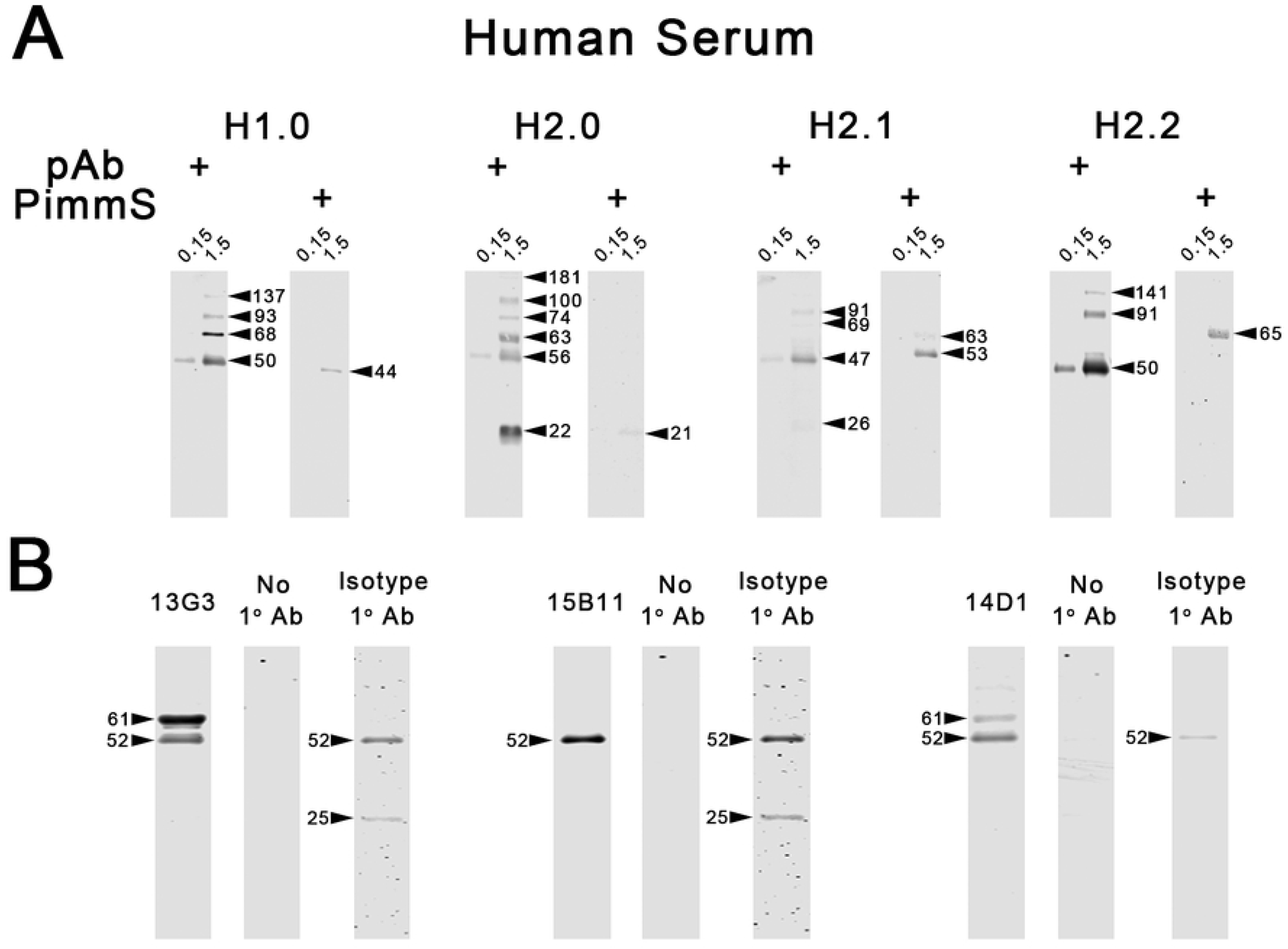
Western blot analyses of CABS1 in human (male) serum. A) In this experiment 0.3 ng/µL for pAbs H1.0 and H2.0 and 0.2 ng/µL for pAbs H2.1 and H2.2 were used and 1.5 and 0.15 µg of serum (n = 1 for each pAb). Preimmune sera (PimmS) controls are shown. B) For mAb 10 ng/µL were used together with 10 µg of serum (n = 3 for each mAb). Controls with no primary mAb or with isotype control mAb are shown.

#### mAbs

For studies of CABS1 immunoreactivity in serum, 1 µg of serum was used and 10 ng/µL of mAb (n=3 for each mAb). When probing serum with 13G3 and 4D1, WB experiments showed a specific immunoreactive band(s) at ∼61 in comparison to isotype controls (Fig 6B). No signal was present when probing membrane only with secondary antibody.

In summary, specific immunoreactive CABS1 was detected in serum at ∼ 61kDa with mAb.

#### Immunohistochemical analysis of human testis and SMG

Because the only previous immunohistochemical studies of CABS1 were done with testes of mice [19] and rats [20] and subsequently with pig [21], we conducted immunohistochemical studies of hCABS1 in human testes for comparison. Thereafter, we investigated hCABS1 expression by immunohistochemistry in human submandibular glands.

#### Testes

Fig 7 A, C and G show the normal morphology of a human testis at increasing higher magnifications stained with hematoxylin and eosin. Seminiferous tubules (ST), interstitial Leydig cells (LC) and, within the tubules, primary spermatogonia (SG), Sertoli cells (SC), developing spermatocytes (SP) and spermatids (S) can be seen (labels). Immunoreactivity using mAb 15B11 detected hCABS1 in LC in the interstitial tissue of the testis (Fig 7 B, D and F), and in SG in ST (Fig 7 D, E, H). In some ST developing spermatids (S) were also positive for CABS1 immunoreactivity (Fig 7 F). Controls done with no primary antibody or with isotype control antibody were negative (Fig 7 F top and lower panel). Monoclonal antibodies 13G3 and 4D1 were also used for immunohistochemistry and no differences in immunoreactivity were noted in comparison to 15B11 for all tissues.

**Figure 7.**
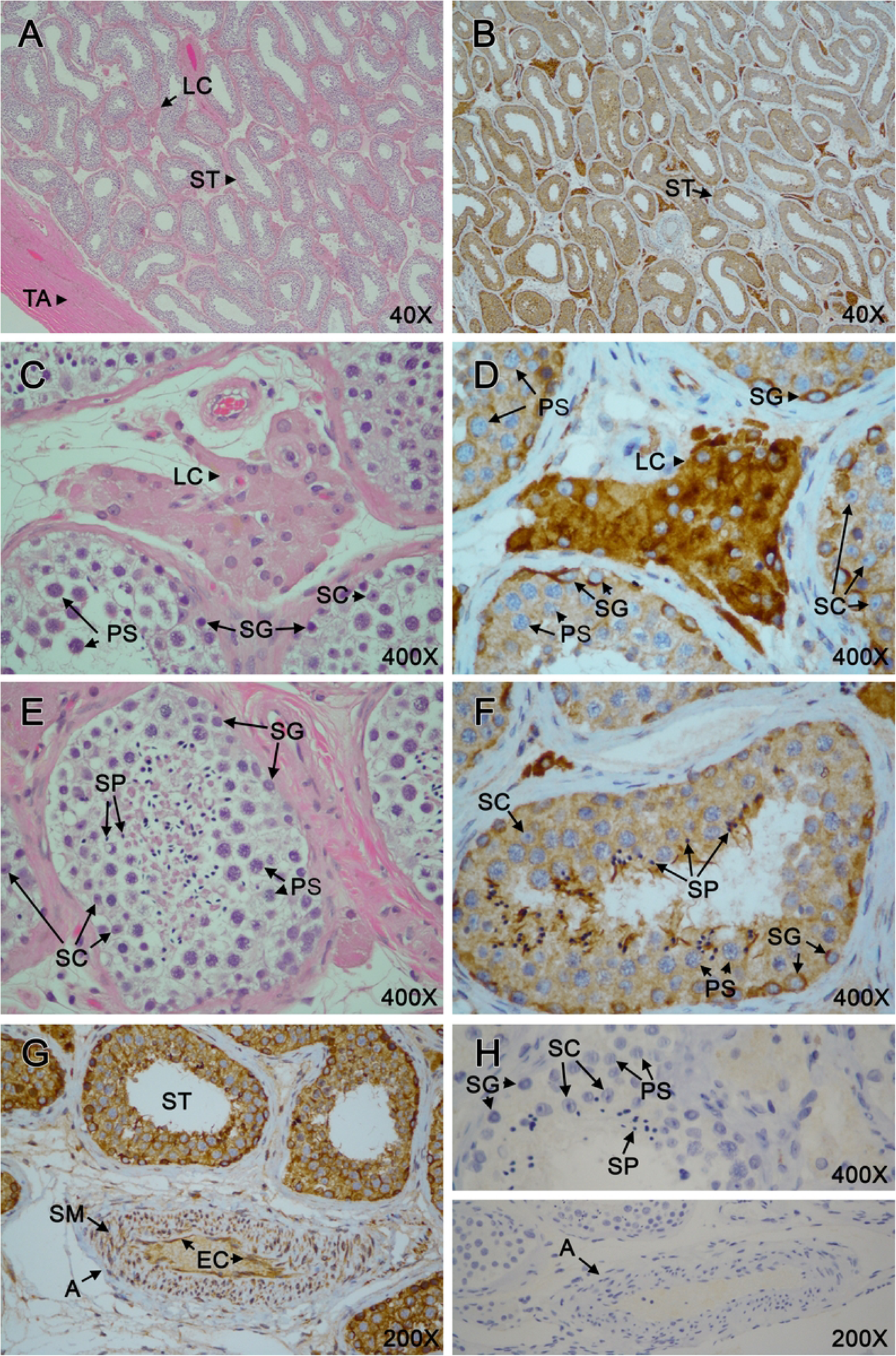
Immunohistochemical analyses of CABS1 in human testes. A) Low power photomicrograph of the connective tissue of the tunica albuginea (TA), seminiferous tubules (ST), and Leydig cells (LC) stained with hematoxylin and eosin (H&E) (40X). B) Low power photomicrograph of seminiferous tubules and Leydig cells stained with 15B11 monoclonal antibody to human CABS1 (40X). C) H&E staining of Leydig cells and, in seminiferous tubules, cell types associated with spermatogenesis (spermatogonia (SG), developing primary spermatocytes (PS) and supporting Sertoli cells (SC) (prominent nucleoli) in 400X magnification. D) Similar image to C, stained with 15B11; note strong staining of Leydig cells, endothelial staining (EC) (400X). E) H&E stain of the different cell types in seminiferous tubule, including spermatids (SP) see also C (400X). F). CABS1 staining observed in seminiferous tubule showing positivity in spermatogonia and spermatids G) CABS1 staining of cells including most intensely stained spermatogonia, spermatocytes and some Sertoli cells; luminal spermatids not prominent. An adjacent artery (A) also exhibits staining in the vascular smooth muscle (SM) and endothelium. H) Negative control (secondary antibody only; isotype controls were also clean) of cells of seminiferous tubules (upper, 400X) and an adjacent artery (lower, 200X).

#### Submandibular glands

Normal morphology of the SMG is shown in Fig 8 A, D at increasing magnification. The lobular structure of the gland can be seen, as well as interlobular vasculature, nerves (N) and excretory ducts (ED). Within the lobes, serous (SA) and mucous acini (MA) are evident, together with numerous sections of smaller salivary ducts (SD) in different plains. Using mAb 15B11, salivary ducts of all sizes had hCABS1 immunoreactivity in epithelial cells (Fig 8 B, E, G, H, J) Controls with no primary antibody or with isotype control antibodies were negative for immunoreactivity throughout the SMG (Fig 8 C, F, I, L). SA cells were positive and the basal, paranuclear regions of MA cells were positive for hCABS1 (Fig 8 E); the mucous-containing regions of these cells were negative for hCABS1. Interestingly, nerves (N) (Fig 8 J, K) were hCABS1 positive and some vessels had hCABS1 immunoreactivity in endothelial cells (Fig 8 K; 8 F shows the negative control for endothelial cells).

**Figure 8.**
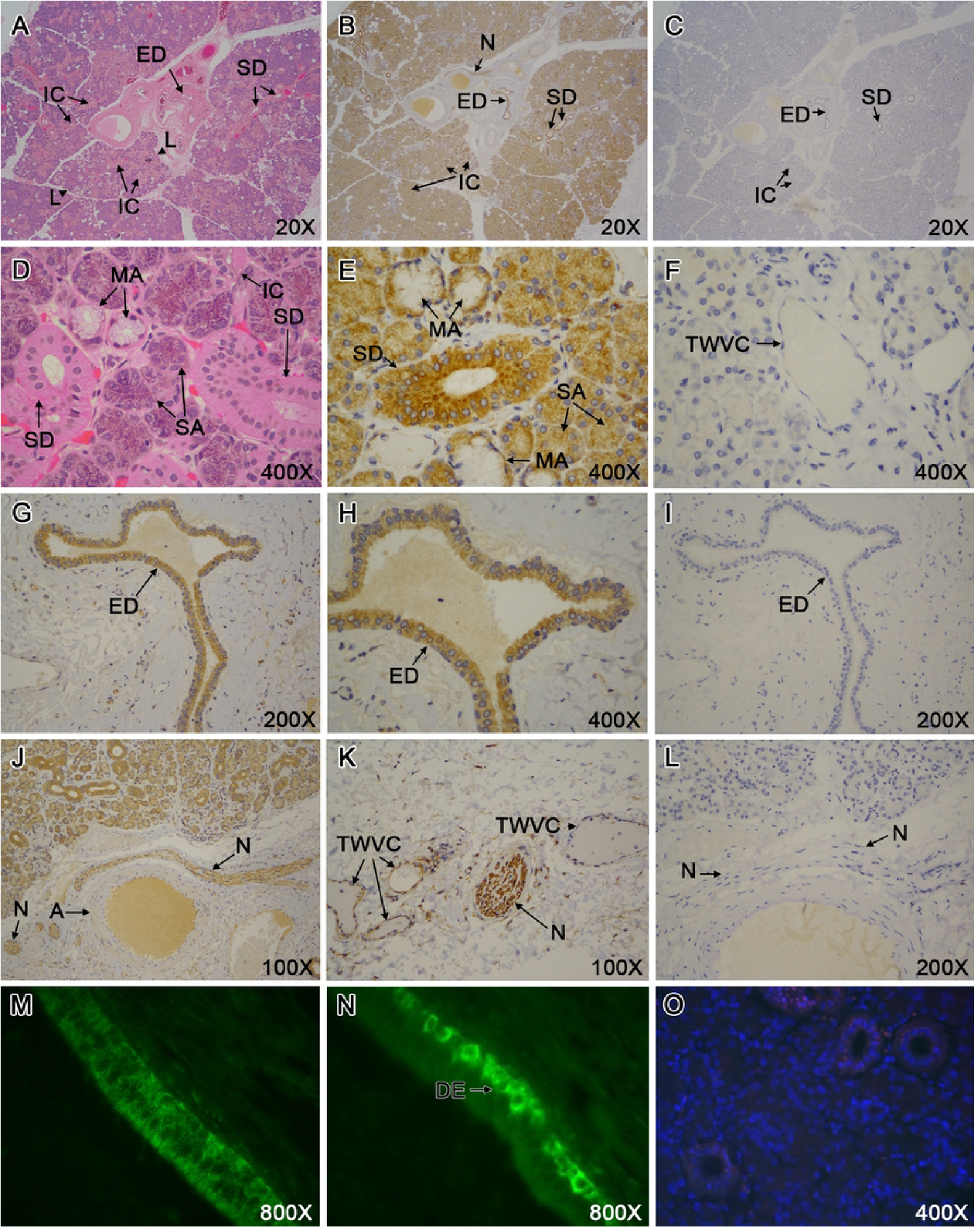
Immunohistochemical analyses of CABS1 in human submandibular glands. A) Hematoxylin and eosin (H&E) staining showing lobes (L) of the gland together with the central excretory salivary duct (ED), intercalated ducts (IC), striated ducts (DD) and associated vasculature (20X). B) Adjacent section showing CABS1 staining of ducts and acini as well as nerve (N) (20X). C) Adjacent section viewed with negative control staining (no primary antibody; isotype controls were also clean) (20X). D) H&E staining of serous (SA) and mucous acini (MA) and associated striated collecting ducts (400X). E) Tissue section viewed at 400X similar to D with CABS1 staining of striated salivary duct and serous and mucous acini (luminal region positive). F) Negative control showing acini and thin wall vascular channel (TWVC), likely a capillary (400X). G) CABS1 staining in excretory duct (200X). H) CABS1 staining in excretory duct showing higher magnification of the ductal epithelium (400X). I) Negative control for CABS1 staining in the excretory duct (200X).J) CABS1 staining in nerves within the salivary gland. Also shown is an artery (A) and another duct (D) (100X). K) CABS1 staining in a nerve as well as in the endothelium of vascular structures in the gland (100X). L) Negative control viewed (no primary antibody, isotype control clean as well) (200X). M and N) Selective immunofluorescent staining of CABS1 (rabbit polyclonal antibodies H2.1 and H2.2, respectively). H2.2 (and H2.0 not shown) showed selective cellular cytoplasmic staining in the basal regions in the epithelium of the excretory ducts (DE), whereas, H2.1 stained the cytoplasm of cells throughout the epithelium. Controls with preimmune sera (e.g. H2.2, [O]) or no primary antibody (not shown) were negative.

Immunofluorescence (see Methods) staining of human submandibular gland sections with pAbs H1 and H2.n (tested independently) yielded a punctate pattern throughout the cytoplasm of all duct cells (Fig 8 M). Interestingly, pAbs H2.0 and H2.2 (tested independently) exhibited CABS1 immunoreactivity in the cytoplasm of specific basal (abluminal) duct cells (Fig 8 N). Analyses of of SMGs immunostained with preimmune sera for the 4 pAb gave no immunoreactivity (Fig 8 O).

#### GEO data assessment of CABS1 tissue distribution and relative abundance

In our search of hCABS1 expression in different tissues in public domain transcriptome databases, we selected three papers that surveyed numerous normal human tissues for gene expression [24–26]. Dezso et al (2008) [25] used the Applied Biosystems human genome microarray of 27,868 genes and 31 normal tissues; She et al (2009) [24] used oligonucleotide arrays from Agilent Technologies Inc, detected 18,149 genes from 42 normal human tissues; Wang et al 2019 [26] used the Illumina HiSeq 2000/2500 system and detected 18,702 protein-encoding genes from 29 normal human tissues. Fig 9 summarizes the normalized relative abundance of hCABS1 from the transcriptomic data from these three studies. CABS1 expression in testes is the best studied of the tissues, but CABS1 mRNA has been found throughout the urogenital, gastrointestinal and respiratory tracts, as well as in glands, the nervous system, immune system and other sites.

**Figure 9.**
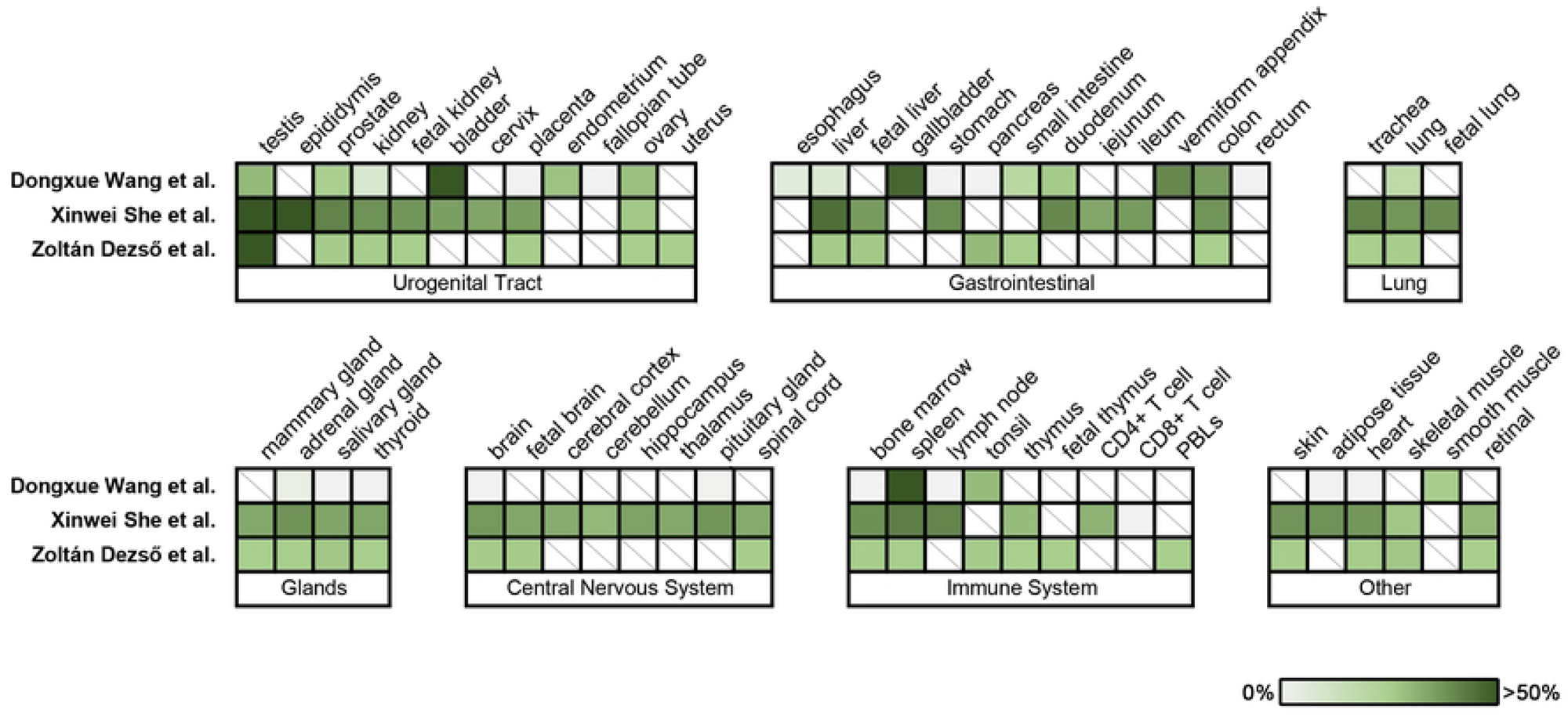
Meta-analysis of the distribution of CABS1 in normal human tissue across three studies. Each study is represented in a row, and tissues are represented in columns. Tissues are grouped in their respective biological system. CABS1 GEO data was extracted and normalized by dividing each expression read by the maximum value observed across tissues within the same study. The normalized percentage of CABS1 expression is shown on a continuous graded scale ranging from 0% (grey) to more than 50% (green), as indicated by the color key. A diagonal line in a box indicates the lack of data within that tissue.

## Discussion

To validate and extend our previous studies with WB [18, 22] and nano-capillary immunoassay (NCIA) [23], herein we described studies of hCABS1 in transient overexpression lysate (OEL), SMG lysates, saliva, serum and testis using pAb and mAb. WB and immunohistochemistry studies were used with Geodata analyses to provide validation for our observations of hCABS1 in these tissues and fluids.

The results of studies with OEL are summarized in Table 1 that shows WB studies with pAb and mAb as well as anti-FLAG antibody. Our previous results using NCIA [23] are also included for comparison (Table 1). Finally, the distribution of CABS1 peptides, determined by mass spectroscopy, in various sections (M_r_ ranges) of a 1 D gel are shown for comparison. M_r_ estimates of 84 and 78 are likely to be the same immunoreactive band given time differences between studies with pAb and mAb, possible technical differences (unknown), and methods of M_r_ calculation. Indeed, results with anti-FLAG antibody confirm that 84 (pAb) = 78 (mAb) (see Fig. 2-B, 4D1).

**Table 1.**
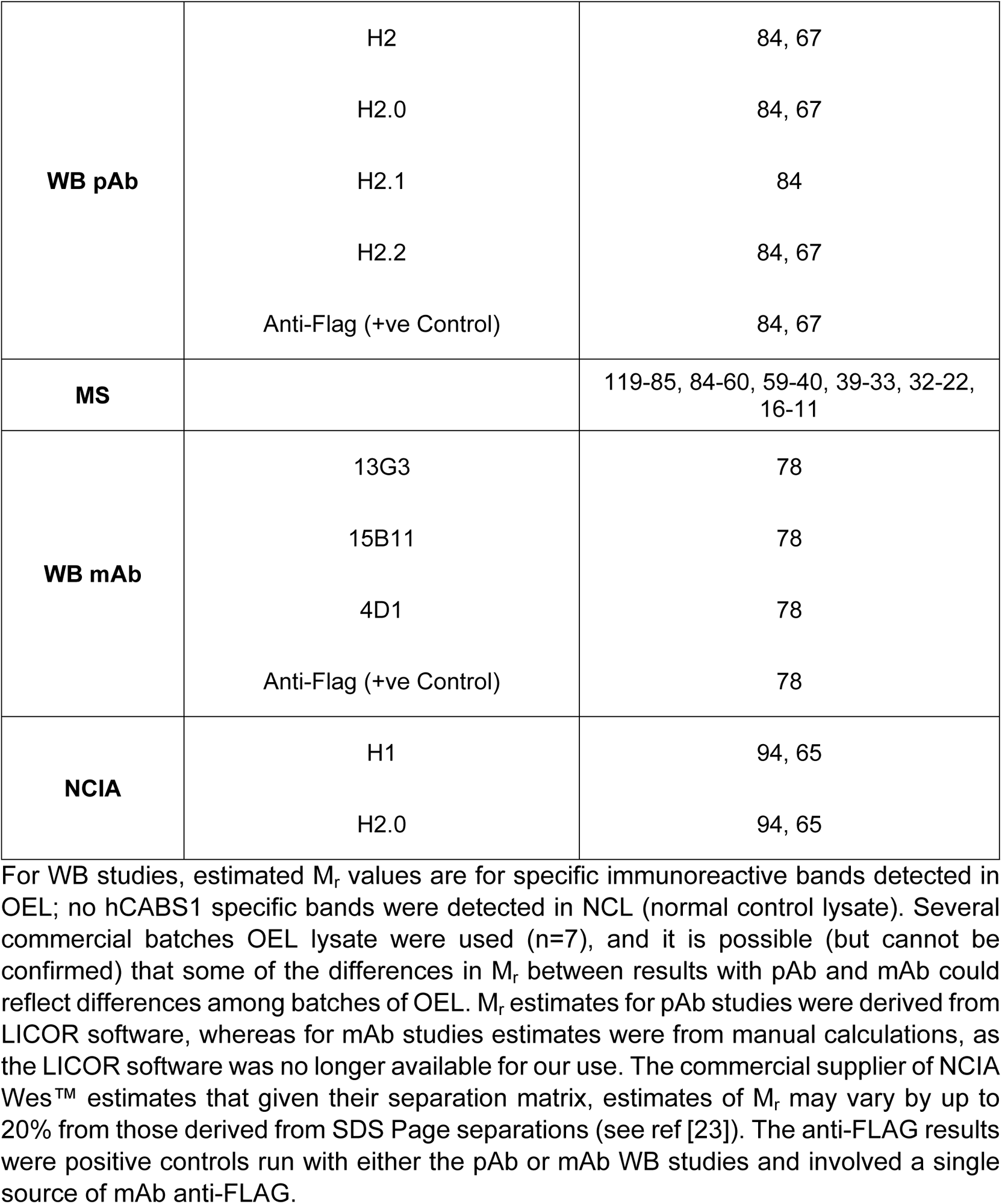
Estimated M_r_ of CABS1 Detected in Transient Overexpression Lysate (OEL) by pAB and mAb in western blot (WB) and Nano-capillary immunoassay (NCIA)

|  |  |  |
| --- | --- | --- |
| **WB pAb** | H2 | 84, 67 |
|  | H2.0 | 84, 67 |
|  | H2.1 | 84 |
|  | H2.2 | 84, 67 |
|  | Anti-Flag (+ve Control) | 84, 67 |
| **MS** |  | 119-85, 84-60, 59-40, 39-33, 32-22, 16-11 |
| **WB mAb** | 13G3 | 78 |
|  | 15B11 | 78 |
|  | 4D1 | 78 |
|  | Anti-Flag (+ve Control) | 78 |
| **NCIA** | H1 | 94, 65 |
|  | H2.0 | 94, 65 |

For WB studies, estimated $M_r$ values are for specific immunoreactive bands detected in OEL; no hCABS1 specific bands were detected in NCL (normal control lysate). Several commercial batches OEL lysate were used (n=7), and it is possible (but cannot be confirmed) that some of the differences in $M_r$ between results with pAb and mAb could reflect differences among batches of OEL. $M_r$ estimates for pAb studies were derived from LICOR software, whereas for mAb studies estimates were from manual calculations, as the LICOR software was no longer available for our use. The commercial supplier of NCIA Wes™ estimates that given their separation matrix, estimates of $M_r$ may vary by up to 20% from those derived from SDS Page separations (see ref [23]). The anti-FLAG results were positive controls run with either the pAb or mAb WB studies and involved a single source of mAb anti-FLAG.

MS shows CABS1 fragments in six segments of the 1 D gel ranging from 119-85 kDa to 16-11 kDa. Given the peptides identified in these different gel segments, there is no evidence of selective fragmentation, i.e., existence of specific portions of CABS1 in distinct gel fragments.

We have previously published our results with pAbs on immunoreactive bands of hCABS1 in SMG lysates [18]. Multiple bands were detected with H2.0 and other pAb, including: bands at 51, 33, 27, and, in some SMG lysates, at 20, 17, 16 and 11 kDa. In the present study with SMG lysate and using H1, 2.0, 2.1, and 2.2 anti-CABS1 pAbs, specific immunoreactive bands were detected with at least three of four pAbs at ∼71, 52 and 27 kDa (Fig 4A; Table 2).

**Table 2.** Specific Human CABS1 Immunoreactivity (Polyclonal and Monoclonal Antibodies) in Submandibular Gland Lysate, Saliva and Serum.

| **Antibody** | **Polyclonal** | **H1** |  | **H2.0** |  | **H2.1** |  | **H2.2** |
| --- | --- | --- | --- | --- | --- | --- | --- | --- |
|  | **Monoclonal** |  | **13G3** |  | **15B11** |  | **4D1** |  |
| **Submandibular Gland Lysate** |  | 52, 27 | 63, 53 | 71, 52, 27 | 53 | 71, 52, 27 | 63. 53 | 71. 52 |
| **Saliva** |  | 60 | 60 | 60 | 60 | N/A | 60 | N/A |
| **NCIA (Saliva)** |  | 60 | N/A | 58, 34 | N/A | N/A | N/A | N/A |
| **Serum** |  | * | 61 | * | - | * | 61 | * |

\*Pilot study (n=1) only; multiple bands, no obvious consistency
Summary of specific CABS1 immunoreactive bands from Figures 3 to 6 (submandibular glands lysate, saliva, and serum).

With at least two of three anti-CABS1 mAbs, immunoreactive bands were found in SMG lysate at 63 and 53 kDa (Fig 4B, Table 2). Human CABS1 immunoreactive bands were detected using pAbs and mAbs in saliva and serum at similar molecular sizes, 60 to 63 and 52 to 53 kDa, results consistent with our earlier report using NCIA [23] (Table 2). Moreover, using mass spectroscopy, hrCABS1 was detected in OEL was detected in at 120-85, 84-60, 59 – 40, 39-22 and 16-11 kDa (Table 1). Two of three mAbs identified a specific immunoreactive band in serum at ∼61 kDa. However, with four pAbs numerous bands were detected and no obvious reproducible profile could be confidently discerned. Interestingly, CABS1 has been detected in human serum using mass spectroscopy, although only a single peptide was identified that covered 3.54% of the protein [27].

With the newly optimized antigen sample and pAb concentrations, in contrast to previous work [18, 22], we only detected a faint specific band at ∼27 kDa in saliva (Table 2). However, a 27 kDa band was detected in SMG lysate, in mass spectroscopy (32-22 kDa gel fragment) and in NCIA (34 kDa with the caveat of the difference in the gel matrices allowing for up to 20% differences in M_r_ estimation between polyacrylamide gels and NCIA, see above).

Our challenge in detecting a 27 kDa band in human saliva using our newly optimized pAb protocol, and mAb, may be explained by low abundance of this protein. Full-column electrophoresis mass spectroscopic analysis of OEL showed that the highest CABS1 expression levels detected by mass spectroscopy were for molecules in the 84-60 kDa range (Fig 3). The second highest detected range was 6 times lower expression levels in the 39-33 kDa range. Although CABS1 peptides were detected at lower molecular weights, the abundance decreased to 19 times lower for the 32-22 kDa range. The concentration of saliva (2.2 µg/well) and pAb (0.3 ng/µL) that yielded highest specificity in WB profiles in our newly optimized protocol was much lower than concentrations we used in previous pAb studies (saliva, 25 µg; pAb 3 ng/µL) [18]. A combination of lower saliva sample concentration, lower pAb and mAb concentrations, and low 27 kDa CABS1 relative abundance could explain our inability to detect this potential form of CABS1 in saliva. It is also possible that our earlier results [18] were incorrect. Thus, we are unable to confirm the presence of a 27 kDa CABS1 form in human saliva, and this raises concerns about whether the 27 kDa molecular form that we reported to be associated with acute stress is indeed hCABS1; a question that will remain pending until new studies are conducted and replicated by others. Regrettably, to date we have been unable to sequence hCABS1 from human SMG and saliva using mass spectroscopy, possibly because of its low abundance proportionate to other proteins. However, our immunohistochemical and geodata studies confirm the presence of CABS1 in human SMG (see below).

A limitation of our studies is that they were conducted over 6 years with several pAb and mAb and with several samples of OEL and NCL (n = 7-8), and SMG lysates (n ∼ 8). For serum and saliva, care was taken to create sample pools that were aliquoted and frozen (-80C), and not repeatedly frozen and thawed. Western blot optimization protocols varied among antibodies and samples, but were established with careful titrations. Moreover, although M_r_ standards were routinely used, the methods for identification of M_r_ of immunoreactive bands varied; initially this was done using the LICOR software, but beginning in 2019 molecular weights were calculated manually, as the LICOR software was no longer available to us. These many issues likely influenced the variability encountered in M_r_ estimates using WB. Furthermore, it was well-recognized that CABS1 is a structurally disordered protein with conformational plasticity [19, 20, 28–30] and an estimated M_r_ (79 kDa, 66kDa respectively) considerably different from the predicted M_r_ of the newly translated polypeptide (42 kDa). This flexible, disordered structure, although potentially of functional significance, may also contribute to the variability in M_r_ that we detected among the tissues and fluids studied. Another component of the variability in M_r_ for hCABS1 may involve the cell of its origin in the tissues (e.g., spermatids, epithelium, nerves, etc.), cell type-specific processing and CABS1 interactions with charged ligands [28].

As previous literature on the cellular localization of CABS1 protein focused on testes in rodents [19, 20] and pigs [21], we conducted the first immunohistochemical cellular localization of CABS1 in human testes (Fig 8). An earlier study had indicated that CABS1 mRNA was abundant in human testes [31]. We found CABS1 in developing spermatids and sperm in human seminiferous tubules (Fig 7). Interestingly, unlike previous rodent and porcine studies, CABS1 was also found in primary spermatogonia, developing spermatocytes, Sertoli cells and in Leydig cells (interstitial cells that produce testosterone and other androgens) in human testes. In mice [19] and rats [20] CABS1 was found in late phases of spermatid development, including in the inner mitochondrial membrane in rats [20] and in the flagellum in mature sperm [19]. In the pig, CABS1 was found in elongated spermatids and sperm, specifically in the principal and end piece of the flagellum, as well as in the acrosome [22]. Interestingly, the shedding of the porcine acrosome (acrosomal reaction) also includes the loss of CABS1, and moreover, the acrosomal reaction can be inhibited by anti-CABS1 antibodies, without an effect on sperm viability [22]. The knockdown of CABS1 in mice disrupted sperm tail structure, induced an abnormality in the flagellum, specifically at the midpiece-principal region junction, and reduced fertility [32]. Taken together this evidence from rat, mouse and pig suggests that CABS1 is involved in spermatogenesis, sperm motility and the acrosomal reaction. Given the presence of CABS1 in human primary spermatogonia, Sertoli cells and Leydig cells, but not these locations in mice, rats and pigs, it seems likely that hCABS1 may play additional roles in male reproduction in humans, perhaps in association with the unique expression of a peptide sequence near the carboxyl terminus in primates^29^ that has been shown to have anti-inflammatory activity [18].

There are few reports of hCABS1 in the human female urogenital tract [33, 34], although as shown in Fig 9, there are numerous reports of CABS1 mRNA. In transcriptomic [24–26] analyses of healthy human tissues, hCABS1 was identified in endometrium, ovary, fallopian tubes, placenta, cervix and urinary bladder. Cerny et al.[33] showed in bovine oviductal epithelial cells, that the levels of CABS1 transcript changed during the estrous cycle. Moreover, Calhoun et al. [34] showed that exposure to the toxin bisphenol A during fetal development altered CABS1 expression in the fetal uterus and suggested that this may influence uterine function later in life. Clearly, much needs to be learned about the roles of CABS1 in reproductive physiology.

Interestingly, our immunohistochemical studies detected hCABS1 in several anatomical compartments and cell types of SMG, including epithelial cells of acini and ducts. Connecting hCABS1 to a potential anti-inflammatory role in humans, it has been reported that in Sjogren’s syndrome, an autoimmune inflammation of the salivary glands, hCABS1 mRNA levels were decreased 5.4-fold in comparison to normal [35]. This reduction in hCABS1 mRNA in Sjogren’s syndrome was confirmed with laser capture microdissection of the acinar and ductal epithelium of minor labial salivary glands [36]. Our observations that nerves and vascular endothelium and smooth muscle in SMG are also hCABS1 positive are novel observations, but supported by Geodata sets that identify hCABS1 in SMG [24–26] and CABS1 in nerves, endothelium and smooth muscle [24–26]. These studies also suggest that hCABS1 is widely distributed in many other tissues including liver, gastrointestinal tract, respiratory tract, neuro, endocrine and immune systems. Given the structural plasticity of CABS1, its widespread tissue distribution and apparent functional diversity, e.g., from spermatogenesis to anti-inflammatory activities, it is imperative to identify the specific cell types that express CABS1 and its functions.

## Acknowledgements

We wish to thank Sarah Canill, Alberta Precision Laboratories, and Yong-Qiu Doughman, Case Western Reserve University for assistance with immunohistochemical studies. Christopher St. Laurent, University of Alberta, assisted with western blot analyses. ER-S received graduate studentships from the Faculties of Graduate Studies and Research, and Medicine and Dentistry, University of Alberta. MM-P received funding from the Natural Sciences and Engineering Research Council, Canada. ADB received funding from AllerGen NCE Inc, Canada.

